# Dosage Matters: A Randomized Controlled Trial of Rehabilitation Dose in the Chronic Phase after Stroke

**DOI:** 10.1101/441253

**Authors:** Carolee Winstein, Bokkyu Kim, Sujin Kim, Nicolas Schweighofer

**Affiliations:** Division of Biokinesiology and Physical Therapy, Herman Ostrow School of Dentistry and Department of Neurology, Keck School of Medicine, University of Southern California, Los Angeles, CA, USA (CW); Division of Biokinesiology and Physical Therapy, Herman Ostrow School of Dentistry, University of Southern California, Los Angeles, CA, USA (NS, CM); Department of Physical Therapy Education, Department of Neurology, SUNY Upstate Medical University, Syracuse, NY, USA (BK); Department of Physical Therapy, Jeonju University, Jeonju, South Korea (SK)

**Author notes:** Corresponding Author: Carolee Winstein, Division of Biokinesiology and Physical Therapy, 1540 Alcazar St. CHP 155, University of Southern California, Health Sciences Campus, Los Angeles, CA. USA.

**Keywords:** rehabilitation, dose-response, recovery, stroke

## Abstract

**Background and Purpose:** For stroke rehabilitation, task-specific training in animal models and human rehabilitation trials is considered important to trigger inherent neuroplasticity, promote motor learning, and functional recovery. Little is known, however, about what constitutes an effective dosage of therapy.

**Methods:** This is a parallel group, four arm, single blind, phase I, randomized control trial of four dosages of upper extremity therapy delivered in an outpatient setting during the chronic phase after stroke. Participants were randomized into groups that varied in total dosage of therapy (i.e., 0, 15, 30, or 60 hours). Seven hundred and four participants were assessed for eligibility, 50 were eligible to enroll, 45 were randomized, 44 participated and 41 completed the study. Planned primary analyses used linear mixed effects regression to model baseline to post-intervention changes in the Motor Activity Log-Quality of Movement rating (MAL_Q_) and the Wolf Motor Function Test (WMFT) time score as a function of therapy dosage. A series of hierarchical models were constructed using the MAL_Q_ and WMFT.

**Results:** We observed a significant dose response curve: the greater the dosage of training, the greater the change in MAL_Q_, with the dose by week slope parameter of 0.0045 (ΔMAL/hour/week; p = 0.0011; 95% CI = [0.0019; 0.0071]). Over the 3 weeks of therapy, this corresponds to a gain of 0.81 in MAL_Q_ for the 60 hour dose.

**Conclusions:** For mild-to-moderately impaired stroke survivors, the dosage of a patient-centered, task specific motor therapy was shown to systematically influence the gain in quality of arm use in the natural environment, but not functional capacity as measured in the laboratory. We highlight the importance of recovery outcomes that capture arm use vs. functional capacity.

**Clinical Trial Registration:** URL: http://www.clinicaltrials.gov. Unique identifier: NCT 01749358

## Introduction

A controversial issue in stroke rehabilitation is the effect of “rehabilitation dose” ^1^. Clinical trials demonstrate that motor therapy delivered in the subacute to chronic phase can effectively increase spontaneous use and function of the affected limb for patients with mild-to-moderate impairments.^2^ Little is known, however, about what constitutes an effective dosage of therapy.^3,4^ Two recent phase II RCTs, the VECTORS trial in the acute setting^5^, and the Dose-Response trial in the chronic setting,^6^ showed that an increase in dosage of task practice did not result in an increase in functional capacity. Given the discrepancy between functional capacity and use,^7^ and the alignment of arm use with patient preferences,^8^ it is important to understand the efficacy of rehabilitation dose on arm use in everyday tasks. With the widespread clinical and research implications, evidence about meaningful outcomes and therapy dosage is particularly needed.^9^

Two important factors in the design of dose-response studies in rehabilitation are: 1) dosage or number of hours of active therapy provided, and 2) behavioral intervention—i.e. the task practice protocol.^9,10^ For this phase I study, we chose four dosages (i.e., active control, low, moderate, high), and a patient-centered behavioral intervention that was especially designed to enhance arm use.^11^

The primary aim is to test the dosage of task-specific practice that is needed to achieve meaningful recovery of the arm and hand in chronic stroke survivors. Here, we report the primary outcome--the dose-response curve for the immediate pre-post intervention effects.

## Materials and Methods

The data that support the findings reported here are available from the corresponding author on reasonable request.

### Study design

This was a parallel group, four arm, single blind, phase I, RCT of four dosages of arm and hand practice administered in the outpatient setting during the chronic phase after stroke. Participants were randomized into four intervention groups (active control, low, moderate, high) that varied, respectively, in total dose of therapy (0, 15, 30, or 60 hours). The PI team (CW, NS) were kept blinded to randomization until study completion. Please see http://stroke.ahajournals.org for a priori power estimates, and details of recruitment and enrollment (Online Supplement). Therapy was provided in three weeklong bouts of four consecutive visits each separated by one month (train-wait-train, Figure 1). All participants signed an informed consent that was approved by the Institutional Review Board of the University of Southern California, Health Sciences Campus, Los Angeles, CA.

**Figure 1:**
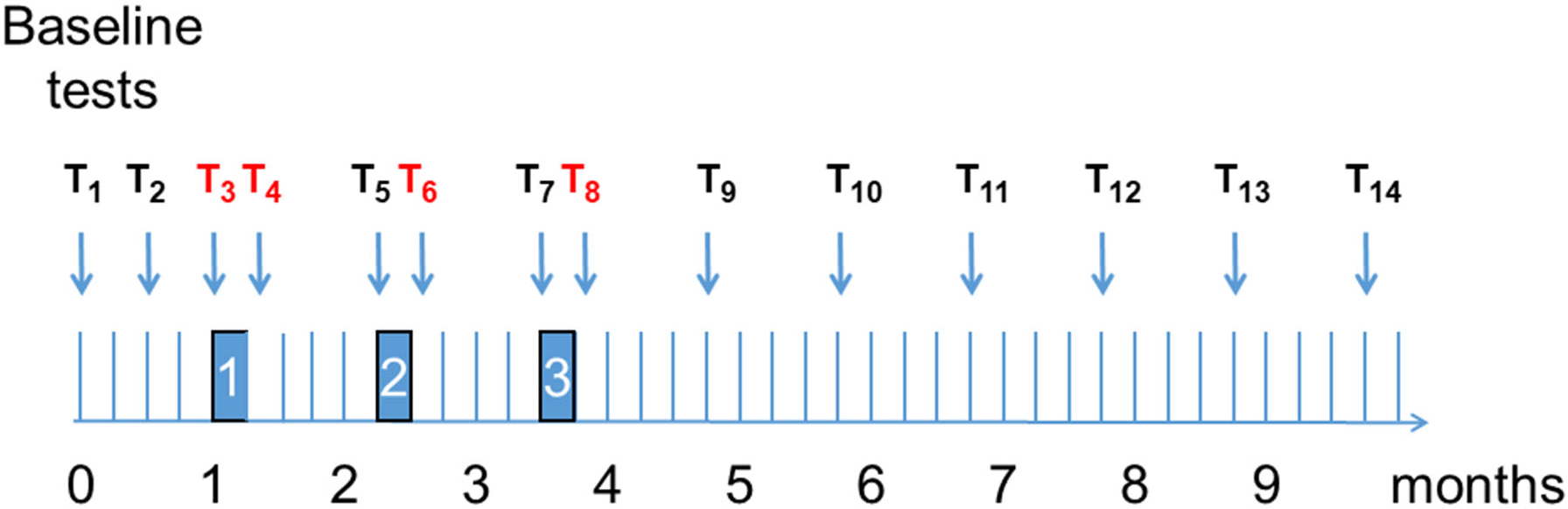
Time-line of train-wait-train training and testing schedule. The total time of the study is 9 months and 3 weeks. The “T”s show the 14 tests. Importantly, the red Ts show the tests used in the dose-response curve analyses described here. The first two “T”s represent the baseline tests (test 1 and 2). The filled rectangles labeled “1”, “2”, and ‘3” show the three 1-week bouts of training. For each week of training, a test was given in the morning of the first day and 3 days after training. Each of the three weeks of training was separated by 4 weeks (“wait periods”). The four groups receive a different number of hours of training within each week of training. All tests included a MAL_Q_ and WMFT assessment. Patients also underwent a structural research grade MRI scan within one month of study enrollment.

### Participants

Participants were screened to ensure they met the inclusion criteria: 1) diagnosis of ischemic or intraparenchymal hemorrhagic stroke without intraventricular extension, 2) stroke onset at least 5 months prior, 3) age ≥21, 4) UE Fugl-Meyer motor and coordination score (UEFM) 19-60/66, and at least a 1 on finger mass extension/grasp release, 5) preserved cognitive function to provide informed consent, and 6) judged medically stable to participate. Participants were excluded if: 1) prior neurologic or orthopedic condition that limited arm/hand use prior to stroke, 2) diminished pre-stroke independence (Barthel Index score < 95, 3) severe arm/hand sensory impairment or neglect, 4) major depressive disorder, 5) severe interfering pain, 6) passive range of motion restrictions: shoulder flexion < 90°, shoulder abduction < 90°, shoulder external rotation < 45°, elbow extension > 20° from full extension, forearm supination-pronation > 45° from neutral, wrist extension < neutral, MCP and IP extension > 30° from full extension, 7) enrolled in rehabilitation or drug intervention, 8) lives too far from training site, 9) received injected or oral anti-spasticity medications, or 10) pregnant.

Seven hundred and four participants were assessed for eligibility; of those, 50 were deemed eligible to enroll, 45 were randomized, and 44 participated in the intervention (See http://stroke.ahajournals.org). After randomization and before intervention, a brain structural MRI scan was performed to characterize the lesion.

Participants were randomized into one of four groups within four strata by severity and chronicity. Severity stratification was based on baseline UEFM score; those with UEFM 41-58 were “mild” and those with 19-40 “moderate”.^12^ Chronicity stratification was based on stroke onset; those 5 months to 1.5 years post-stroke were “early” and those >1.5 years were “late”.

### Therapy Intervention

The intervention was based on the Accelerated Skill Acquisition Program (ASAP), described in detail elsewhere.^11,13^ In essence, ASAP is a personalized program of task-oriented training, that incorporates elements of skill acquisition, and capacity building (i.e., impairment mitigation), with intrinsic motivational enhancements (i.e., patient empowerment). Participant-selected tasks provide the context for practice. ASAP is especially designed to enhance arm use and to foster transfer of meaningful skills from clinic to home/community environment. Importantly, task practice, favors development of high-quality, skilled movements rather than high volumes of repetitions. Therapy was delivered by three physical therapists trained and standardized according to the foundational principles of ASAP.

### Recovery outcomes evaluation

Primary outcome measures included the MAL_Q_ rating and WMFT time score. The MAL_Q_ is a valid and reliable patient-reported outcome measure, consisting of a semi-structured interview in which participants recall and rate the quality of movement of the paretic arm for 28 activities of daily living performed outside the laboratory.^14^ The laboratory-based WMFT time score is a valid and reliable measure of motor performance for 15 hierarchically arranged arm and hand goal-directed tasks.^15^ Each of these assessments were administered bi-weekly in the month before training (baseline Test 1, 2), and, for each of the three 1-week training bouts, the morning of training day 1 and 3 days following each training bout (Figure 1). Trained and standardized research assistants, blinded to group assignment, performed all assessments.

### Statistical Analysis

For planned primary analyses, we used linear mixed effects regression (LMEs) to model changes in MAL_Q_ and WMFT during treatment as a function of dosage group (0, 15, 30, and 60 hours), similar to Lang and colleagues.^6^ Compared to repeated-measure ANOVA, LME is the proper choice of statistical method, notably because LMEs allow: 1) flexible modeling of individual trajectories over time, thereby minimizing the effects of measurement noise, 2) model building and comparison that uses time as a continuous variable or a categorical variable, 3) preservation of data from all participants for whom one or more assessment point may be missing, and 4) to capture the high variability in lesion, impairment, spontaneous recovery, and responsiveness to therapy post-stroke.^16^

For each of the primary outcomes (MAL_Q_ and WMFT), we developed two types of dose response models in which dose was treated either 1) as a continuous variable or 2) as a categorical variable. Because training was given in three 1-week bouts, each spaced by one month, we studied the effect of training by concatenating the three training bouts (therefore using Tests 3, 4, 6, and 8, see Figure 1). T<u>he test of a significant dose response relationship is given by the significance of the fixed effect coefficient of the interaction term, *dose x week*, in the continuous model.</u> The coefficients are in units of MAL_Q_ and WMFT (log transformed) change per week, per hour for continuous models, and in units of MAL_Q_ and WMFT change per week for categorical models.

In secondary analyses, we consider three co-variates in model development based on previous research: age, ^17,18^ chronicity, ^19^ and concordance. ^6^ We also tested for effects of baseline for each dependent variable (using the average of MAL_Q_ or WMFT from baseline Tests 1 and 2). These variables were entered as modifiers, intercepts and slopes.

For nested models, choice of co-variates, random effects, and random-effects covariance structures were based on the Log-likelihood ratio test. For non-nested models, comparison was based on minimum Bayesian Information Criteria (BIC), which provides a measure of quality of fit by minimizing fitting error and penalizing the number of model parameters. Residuals were examined for normality and the presence of outliers. Statistical analyses were performed with R (R Foundation for Statistical Computing, Vienna, Austria), with the *lmer* functions of the *lmerTest* library. Statistical significance threshold was set at p < 0.05.

Finally, to verify our assumption that concatenating the three 1-week training bouts does indeed represent a true dose response relationship, we confirmed that there was no overall change in primary outcomes between bouts. For this, we used similar models developed to test a continuous dose relationship but tested for changes between Tests 4 and 5, and changes between Tests 6 and 7 (Figure 1). For both testing periods, and for both outcomes, the fixed effect parameter of *week* and *dose x week* was not significant. The dynamics of the response, including the train-wait-train intervention, and follow-up period are the focus of a secondary outcome analysis.

## Results

### Participants

Data from 41 participants who completed the study were included. Demographic and clinical profiles are presented in Table 1. Generally, there were no baseline group differences in non-study relevant demographic and clinical characteristics, with the exception of MAL_Q_. Participants were evenly allocated into groups with respect to severity (i.e., lesion overlap, motor impairment) and chronicity (i.e., early or late). No sex-based or racial ethnic-based differences were present (sex, p = 0.68, ethnicity, p = 0.34, race, p = 0.32, data not shown).

**Table 1.**
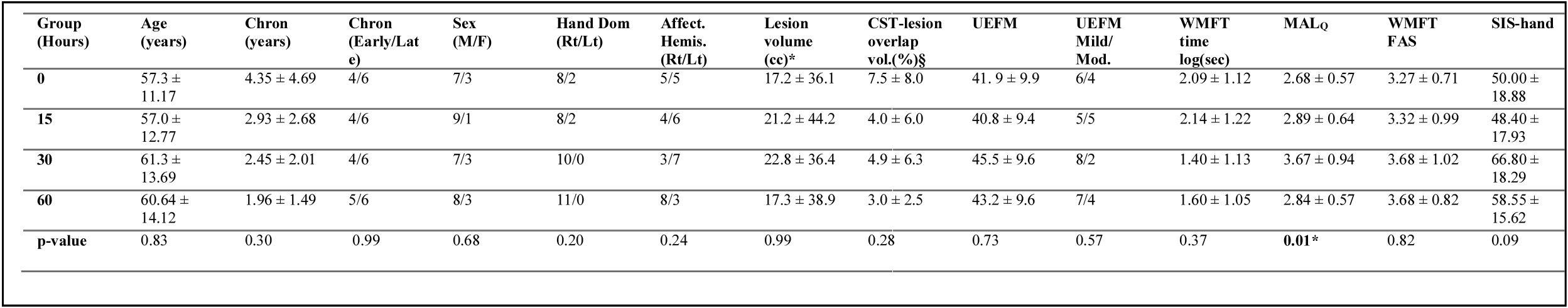
Baseline Demographic and Clinical Characteristics Information including Lesion Characteristics. Chron = Chronicity. Dom = Dominance, Affect Hemis = Affected Hemisphere, UEFM = upper extremity Fugl-Meyer, WMFT =Wolf Motor Function Test, MAL_Q_ = Motor Activity Log-Quality of Movement, WMFT FAS = WMFT Functional Ability Score, SIS-hand = Stroke Impact Score-Hand Domain. §Note 1: Binary lesion masks were drawn on each participant’s T1-weighted images. The number of lesion voxels were counted, and lesion volumes were calculated in cubic centimeter (cc). †Note 2: 3-D template CST (cortico-spinal tract) images were transformed to each participant’s T1-weighted image space. We counted the number of overlap voxels between binary template CST mask and lesion mask. We calculated the proportion of CST-lesion overlap volume to the entire CST volume. *Note 3: The locus of significant group difference in Baseline MAL_Q_ is the 30-hour group. Notice that the 30-hour group also exhibited a higher Baseline SIS-hand score, slightly higher UEFM score, and slightly faster WMFT time score, compared to other dose groups; though these latter differences were not reliable as the ANOVA analyses demonstrate.

Lesion analysis (See http://stroke.ahajournals.org) shows that on average, 4.9 ± 6.0 % of ipsilesional CST was compromised (median: 2.8 %, 25-75 percentile: 0.6% - 5.4%), indicating mild-to-moderate injury to the descending motor pathways (Table 1). ^20,21,22^

### Dose-response: primary analyses

#### *Primary outcome*—MAL_Q_

The final models using the change in MAL_Q_ for both continuous and categorical dose was (in Wilkinson notation):

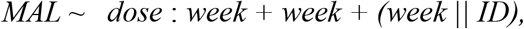

where the operator “:” represents interactions, ID is the subject, (*week ║ID*) represents both random slopes and intercepts; and the operator ║ indicates a diagonal random effect covariance structure, which provides a better model than the full covariance structure. Figure 2 shows the individual data (dot) and superimposed continuous dose mixed effect model with random effects (lines); note how this simple dose-response model, with only three fixed effect parameters (see Table 2; left column) provides an overall excellent fit to the data.

**Figure 2:**
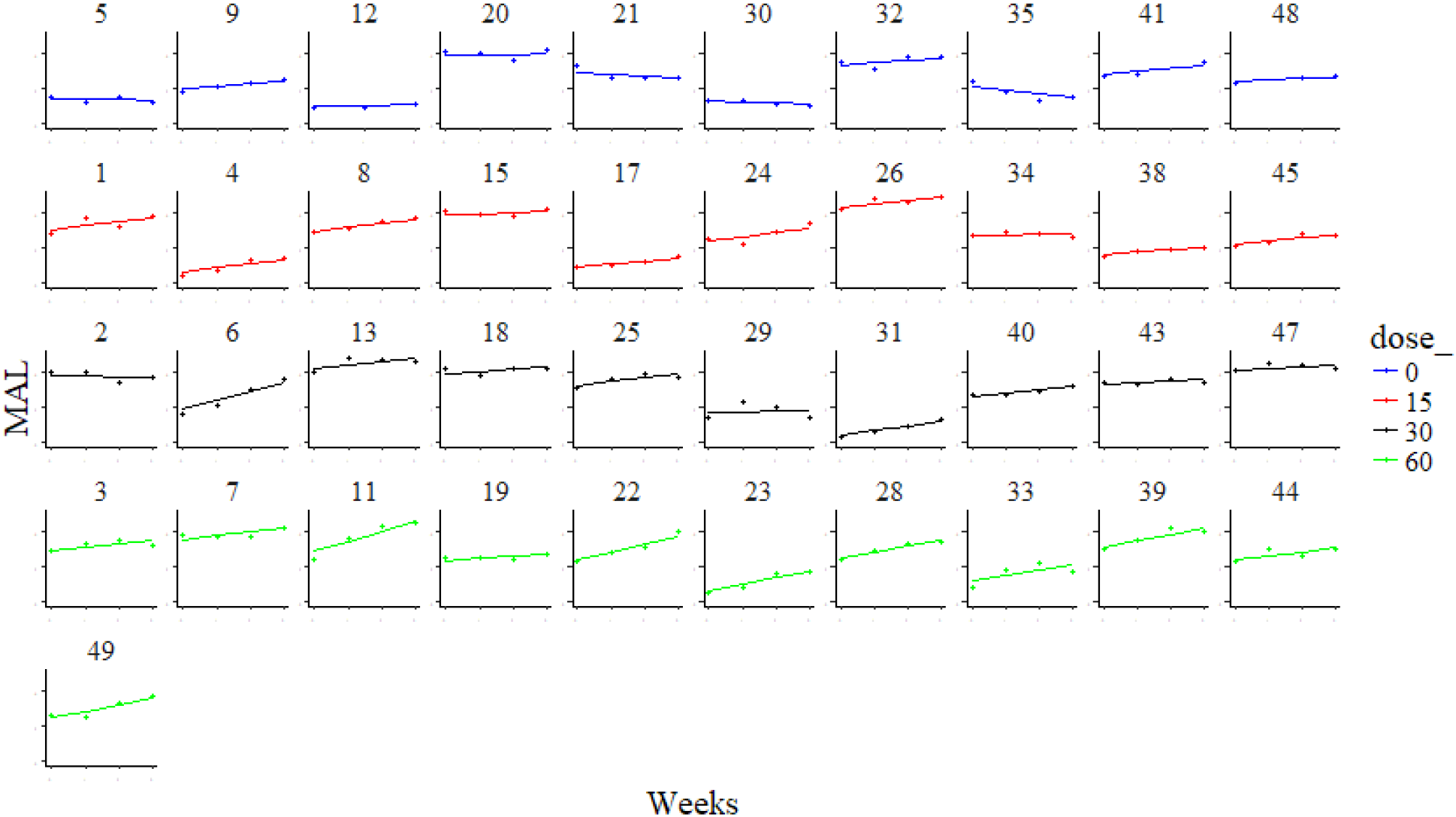
Individual participant plots showing MAL_Q_ change by test week. Each row illustrates group data and model fit for: blue = 0, red = 15-hour, black = 30-hour, green = 60-hour. ID numbers correspond to a subject. Individual participant data (dot) and super-imposed continuous dose mixed effect model (lines) for MAL_Q_. Participant data were recorded before the start of the intervention period and after each week of training or active control. Note that a single mixed model was plotted for all dosages and all participants. Notice the excellent fit, and the larger slopes for many participants in the 60-hour group (two bottom rows).

**Table 2:** Change in MAL_Q_ per dose and per week. Fixed effect coefficients from continuous and categorical models from the dose-response analyses for the primary outcome variable MAL_Q_. Note how the interaction coefficient, *dose_cont:week*, which defines the dose-response, is significant. MAL_Q_ = Motor Activity Log-Quality of Movement.

|  | *Dependent variable:* |  |
| --- | --- | --- |
|  | MAL |  |
|  | Continuous dose | Categorical dose |
| week | 0.052<br>(0.046) | 0.008<br>(0.058) |
| dose_cont:week | 0.005***<br>(0.001) |  |
| dose_15:week |  | 0.185*<br>(0.081) |
| dose_30:week |  | 0.153<br>(0.081) |
| dose_60:week |  | 0.311***<br>(0.079) |
| Constant | 2.542***<br>(0.168) | 2.542***<br>(0.168) |
| Observations | 164 | 161 |
| Log Likelihood | -114.335 | -113.130 |
| Akaike Inf. Crit. | 240.669 | 242.261 |
| Bayesian Inf. Crit. | 259.158 | 266.912 |
| *Note:* | *p<0.05; **p<0.01; ***p<0.001 |  |

The continuous dose MAL_Q_ model reveals a significant dose response curve (Table 2; left column): the greater the dosage of training, the greater the change in MAL_Q_, with the dose by week (*dose_cont:week* in Table 2) parameter of 0.0045 (ΔMAL/hour/week; p = 0.0011; 95% CI = [0.0019 - 0.0071]). Given the slope, the corresponding mean total MAL_Q_ increase for 60 hours was 0.27 per week on average, and therefore 0.81 in three weeks (Figure 3). In contrast, the effect of *week* was not significantly different from zero, indicating that there was no testing effect (p = 0.26). In addition, note that the simple *dose* regressor was not included as an effect in the final model, indicating that the groups were relatively well balanced for MAL_Q_ (despite larger baseline MAL_Q_ for the 30-hour group, Table 1). The corresponding dose-response curve over the three weeks of training is shown by the red line in Figure 3.

**Figure 3:**
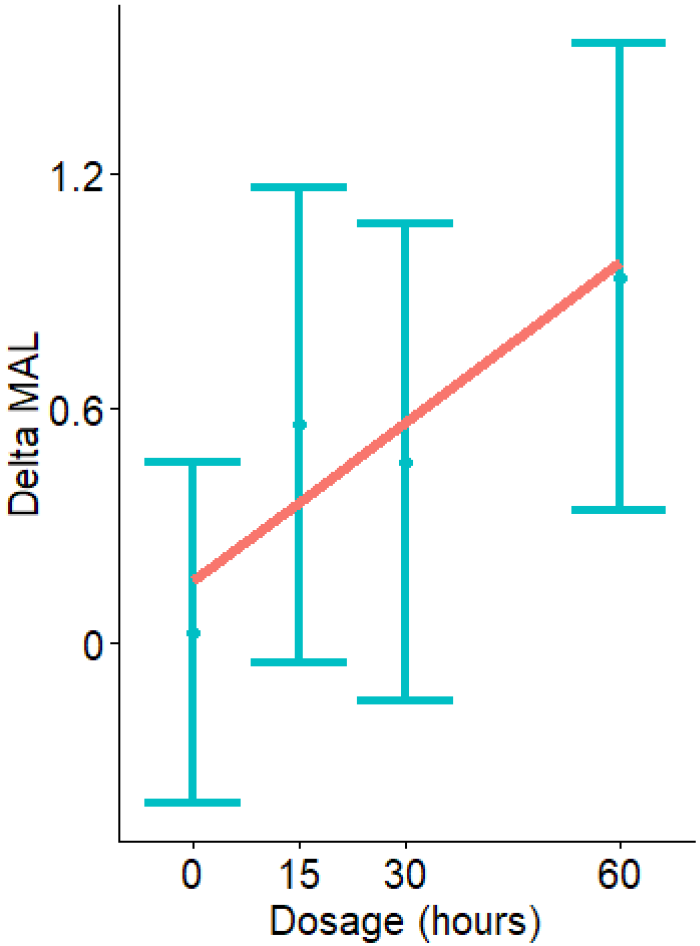
Change in MAL_Q_ over the intervention period as a function of dosage. The red line shows the significant dose response curve (p = 0.0011) derived from the continuous model. The error bars show fixed effect coefficients for dose-response from the categorical model.

The categorical dose MAL_Q_ model (Table 2; right column) shows that the dose response is largely driven by changes in MAL_Q_ for the 60-hour group, as the *dose_60:week* coefficient is significantly different from the 0 dose *week* parameter (mean ± SE: 0.31 ±0.079; p = 0.00031). The mean total MAL_Q_ increase over 3 weeks for a 60 hour dose compared to the active control corresponds to 0.93 (Figure 3). In addition, the *dose_15:week* coefficient is also significantly different from the 0 dose *week* parameter (i.e. the constant in Table 2; p = 0.027). Comparison of the continuous dose model to the categorical model in Figure 3 shows that the change in MAL_Q_ as a function of dose is approximately linear, with a less-than-linear increase for the 30-hour dose, however.

#### Primary outcome— WMFT (sum of time; log transformed)

The final model using the WMFT (log transformed) for both continuous and categorical doses was:

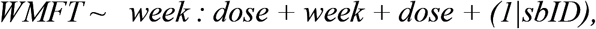

where *(1|sbID)* is the random intercept. The continuous dose model reveals no significant dose response curve for the WMFT (Table 3). The *dose x week* interaction parameter was not significant (p = 0.60), showing no differences in the change of WMFT time score between dosages. The effect of time (week) in the active control group was not significantly different from zero, although approaching significance (-0.045, p = 0.057). Similarly, the categorical model shows that no dose of ASAP improved WMFT compared to the active control group, as none of the categorical *dose x week* parameters were significant.

**Table 3:** Change in WMFT (log) per dose and per week. Fixed effect coefficients from continuous and categorical models from the dose-response analyses for the primary outcome variable log of the WMFT time-response. WMFT = Wolf Motor Function test.

|  | *Dependent variable:* |  |
| --- | --- | --- |
|  | WMFT |  |
|  | Continuous dose | Categorical dose |
| dose_cont | -0.009<br>(0.007) |  |
| dose_15 |  | 0.058<br>(0.444) |
| dose_30 |  | -0.682<br>(0.444) |
| dose_60 |  | -0.435<br>(0.434) |
| week | -0.045<br>(0.023) | -0.038<br>(0.031) |
| dose_cont:week | -0.0003<br>(0.001) |  |
| dose_15:week |  | -0.043<br>(0.042) |
| dose_30:week |  | 0.015<br>(0.042) |
| dose_60:week |  | -0.036<br>(0.041) |
| Constant | 2.012***<br>(0.247) | 2.041***<br>(0.314) |
| Observations | 161 | 161 |
| Log Likelihood | -69.102 | -67.083 |
| Akaike Inf. Crit. | 150.204 | 154.166 |
| Bayesian Inf. Crit. | 168.693 | 184.980 |
| *Note:* | *p<0.05; **p<0.01; ***p<0.001 |  |

#### Dose-response: secondary analyses

In secondary analyses models, we included age, chronicity, concordance, and the corresponding baseline outcome measures. For neither MAL_Q_ nor WMFT models, were age, chronicity, or concordance significant modifiers of either the intercept or dose response-curve (all p > 0.1).

## Discussion

Our primary outcome findings demonstrate that the higher the dosage of ASAP, the greater the change in Motor Activity Log-Quality of Movement over three spaced training bouts, (~4 months) in chronic stroke survivors. The average magnitude of overall change in MAL_Q_ for the 60 hour group (i.e., 0.81) is clinically meaningful, particularly given the chronicity of this cohort.^23^ This is the first reported dose-response effect for motor therapy in chronic stroke survivors with mild-to-moderate motor impairment. However, only one of the two primary outcomes were responsive to dosage. Dosage modified the participation-level outcome, a measure of arm and hand use, but not the activity-level outcome, a measure of functional capacity.

There is precedence for dose-response insensitivity using activity/functional capacity outcomes. The recent phase II RCT sought to determine the dose-response of task specific arm/hand training in people at least 6 months after stroke.^6^ A large number of practice trials were administered, but the investigators found little evidence that dose/number of repetitions or active therapy time made a difference. Thus, neither our study nor Lang et al’s study^6^ observed a dose-response effect for a laboratory-based measure of activity/functional capacity.

Considerable controversy prevails in the clinical research community as to what constitutes a meaningful outcome measure in the context of human clinical trials in rehabilitation.^10,24,25,26^ Recent efforts to develop consensus in the field are underway.^27^ Importantly, for this phase I RCT, we were particularly interested in how distinct aspects of recovery are affected by dosage. Most task-specific training programs are designed to target a specific construct of importance for recovery.^28^ In secondary analyses, Lang and colleagues examined participation level outcomes using the SIS-hand^29^ and COPM^30^ (performance and satisfaction), but there again, found no effect of therapy dosage.^6^ Interestingly, 90% of their participants perceived a meaningful change, attributed to therapy; this perception, however did not correspond to an ARAT change score. By design, ASAP targets skilled (high quality) paretic arm use (participant-chosen use) in the natural environment.^11,31^ Theoretically, it is through continued use in the natural environment that the associated gains in functional capacity are more likely to emerge, long-term.^32,33^ From this perspective, it is not surprising here, that the immediate effects of dosage were evidenced by gains in arm and hand use, outside the laboratory. ASAP emphasizes skilled arm use in the natural environment rather than artificial task demonstration performed “as fast as possible” (i.e. as with WMFT).

An alternative and complementary explanation for the discrepancy in findings between our two primary outcomes is the possibility that ASAP therapy effectively reduced non-use behavior directly through its collaborative, patient-centered approach, but had only indirect, and perhaps lesser effects on functional capacity. This idea is consistent with results of the ICARE RCT to determine the efficacy of ASAP sub-acutely after stroke.^34,24^

### Limitations

Our study has limitations. While there was a dose response for quality of arm use, the 30-hour group may have benefited somewhat less from the intervention because relative to the three other dosage groups, baseline MAL_Q_ was higher. Further, the 30-hour group was less disabled (though not statistically so) on other baseline metrics (i.e., SIS hand, UEFM, and WMFT), perhaps warranting a smaller response to dosage. Because of the relatively small cohort, we chose to limit number of stratification factors in controlling baseline differences. Taken together, in spite of these limitations, and a relatively small sample, we provided phase I evidence that dosage of motor therapy does matter for mild-to-moderately impaired chronic stroke survivors when the recovery outcome is arm use.

### Summary/Conclusions

For the first time, we provide evidence for a significant dose-response relationship for a motor therapy delivered in the chronic stage after stroke. Patients with mild-to-moderate motor impairment showed a meaningful change for a participation-level outcome but not for an activity/functional capacity-level outcome with a higher dosage of task-specific training. We highlight the importance of recovery outcomes that reflect arm use vs. functional capacity.

## Acknowledgements

None

